# Antigenic landscape of a highly mutated SARS-CoV-2 Spike in ongoing viral evolution

**DOI:** 10.64898/2026.02.27.708159

**Authors:** Priscilla Turelli, Elise Eray, Charlène Raclot, Didier Trono, Aleksandar Antanasijevic

## Abstract

The ongoing emergence of SARS-CoV-2 variants in an increasingly immune-experienced population is largely enabled by the plasticity of the Spike protein, which lies at the frontline of host immune pressure. Here, we investigated how extensive remodeling of the N-terminal domain (NTD) in Spike influences its antigenic properties. Using BA.2.87.1, a variant heavily mutated in this domain, we found that even large deletions do not substantially disrupt overall Spike structure or pseudovirus infectivity. However, our structural and binding analyses revealed that the NTD exhibits increased flexibility and that interaction with a heme-metabolite, contributing to assembly of an immunodominant epitope, is lost. These conformational effectscoupled with mutations, compromised the ability of human convalescent antibodies to engage this domain, contributing to their reduced neutralizing capacity. Consequently, BA.2.87.1-like variants may escape recognition by the pre-existing NTD-targeting antibodies, potentially reducing protection. Together, our results highlight the intrinsic adaptability of the Spike beyond the receptor-binding domain, with important implications for immune escape during viral evolution.

**Teaser:** Structural flexibility of the Spike and immune escape of a SARS-CoV-2 variant with public health implications.

## Introduction

After acquiring early post-zoonotic mutations that improved adaptation to humans, SARS-CoV-2 has continued to evolve under the combined pressures of viral fitness and host immunity in a population that is now largely vaccinated (>70%) and previously infected (>779 million reported cases by January 2026) (*1*). The Spike glycoprotein, exposed on the viral surface, mediates attachment to the angiotensin-converting enzyme 2 (ACE2) receptor (*2*–*5*) and subsequent conformational changes leading to viral entry. Its surface accessibility and central role in host-cell entry make Spike the primary target of neutralizing antibodies (nAbs), driving its adaptative evolution. The major nAb epitopes are located in the receptor binding domain (RBD) (*6*, *7*) and N-terminal domain (NTD) of the Spike S1 subunit (*8*, *9*), as well as in the stem helix (SH) and fusion peptide (FP) of the S2 subunit (*6*). Since ACE2 engagement is critical for viral entry, the bulk of neutralization in human sera predominantly arises from RBD-specific antibodies, with NTD-specific antibodies providing a smaller contribution (*10*). As such, these domains face strong B-cell immune pressure, and while the RBD is constrained to preserve ACE2 engagement, the NTD shows greater sequence variability consistent with enhanced antigenic plasticity (*11*).

Some potent NTD-targeting nAbs are public antibodies (*12*, *13*) that bind a common antigenic supersite spanning residues Q14-T20, F140-R158 and H245-A264 in the NTD loops N1, N3 and N5, respectively (*13*). The conformation of these domains and presentation of underlying epitopes partially depend on the binding of heme metabolites into the corresponding heme-binding pocket of the Spike (*14*, *15*). Since the pandemic began, insertions and deletions in this immunodominant domain have repeatedly arisen in most circulating variants, including N3:Δ144 (Alpha), N3:Δ156-157 (Delta), N5:Δ242-244 (Beta) and N5:Δ247-253 (Lambda). Consistently, deletions in N1 and N3 loops and insertions in N1 are found in Omicron variants, altogether indicating a continued and persistent selective pressure on this domain. Structural and functional studies have shown that such mutations remodel the NTD surface, often abrogating binding by NTD-specific nAbs. Nevertheless, the full extent of NTD flexibility and its impact on immune evasion remain incompletely characterized, limiting our ability to anticipate viral evolution and guide future vaccine and therapeutic design efforts.

The BA.2.87.1 isolate is an extreme example in which the entire N1 loop and 37% of N3 loop of the NTD supersite are deleted. This lineage was first identified in South Africa at the end of 2023, with the first sequence deposited in January 2024 (EPI_ISL_18845398). BA.2.87.1 carries an exceptionally high total of 65 amino acid mutations in the Spike protein relative to the original Wuhan reference strain, 33 of which map to the NTD. Because of its extensive mutations, including deletions, and its potential alteration of immunogenicity compared with previous circulating variants, it was rapidly classified as Variant Under Monitoring by the European Center for Disease Prevention and Control. Previous studies have shown that BA.2.87.1 can evade certain monoclonal nAbs and serum antibodies induced by infection and/or vaccination more efficiently than the ancestral B.1 or its BA.2 parental strain, but less efficiently than its contemporary JN.1 lineage (*16*–*21*).

In this study, we build on prior work and use BA.2.87.1 as a model system to interrogate how extensive sequence-level remodeling reshapes the structural, functional, and antigenic properties of the SARS-CoV-2 Spike protein. By integrating high-resolution cryo-electron microscopy (cryo-EM), pseudovirus infectivity assays, and surface plasmon resonance, we show that the NTD can simultaneously accommodate large deletions, amino-acid substitutions, glycan remodeling, and loss of heme binding while preserving overall structural assembly and maintaining infectivity comparable to other circulating strains. Because these mutations map to surface-exposed regions of the NTD, they drive pronounced antigenic changes, resulting in a loss of detectable NTD-directed binding by pre-existing SARS-CoV-2 antibodies and lower neutralization. Together, our findings highlight the NTD’s structural robustness coupled with extreme antigenic flexibility and raise concern that continued Spike evolution could extend this flexibility to other regions, potentially producing variants with minimal cross-reactivity to pre-existing SARS-CoV-2 antibodies.

## Results

### BA.2.87.1 infectivity is lower but comparable to its contemporary variants

Small insertions and deletions in the NTD have been observed recurrently in past circulating variants (**Fig. 1, A and B**). While the small deletion at positions H69-V70 has been associated with enhanced viral infectivity (*22*), larger deletions in the NTD are generally expected to reduce infectivity. Considering the potentially unpredictable epistatic effects of the numerous and often unique mutations in BA.2.87.1 Spike (**Fig. 1, B and C**), we first compared the infectivity of virus-like particles (VLPs) pseudotyped with BA.2.87.1 Spike to those carrying the Wuhan Spike or 21 known successful Omicron sublineages Spikes (**Fig. 1, D and E and fig. S1 and S2**). We used SARS-Cov2-based VLPs (SC2-VLPs) (*23*) as a surrogate system, which better recapitulate key features of authentic SARS-CoV-2 compared to non-native models based on different viruses with a fundamentally different assembly process. Since the Spike is processed by cellular proteases (e.g., furin at the S1/S2 site and TMPRSS2 at the S2’ site) and cleavage efficiency has been linked to increased infectivity (*24*, *25*), we examined both the cleaved and uncleaved forms of the protein in producer cells and VLPs. Overall, we observed similar ratios of cleaved to full-length Spike proteins across all producer cells, with the BA.2.87.1 Spike showing proteolytic processing comparable to its parental BA.2 Spike and slightly lower cleavage efficiency than its contemporary JN.1 Spike (**Fig. 1D**, upper panel). The uncleaved form was, as expected, more difficult to detect in VLPs purified by ultracentrifugation over a sucrose cushion, but the ratio of S2 versus S2’ cleaved products was lower for BA.2.87.1 compared to BA.2 (**Fig. 1D**, lower panel). Because BA.2.87.1 efficiently utilizes TMPRSS2 (**Fig.1D** and (*21*)), we engineered an ACE2+/TMPRSS2+ HEK293T stable cell line to assess SC2-VLPs infectivity. As BA.2 was the parental strain for BA.2.87.1 and for successful sublineages still dominating worldwide, including JN.1 and XBB descendants, we used SC2-VLPs coated with BA.2 Spike as a reference for comparison. When normalized for particle number, we observed overall similar infectivity among the 23 different VLPs tested (**Fig. 1E**). We recorded a few exceptions where VLPs carrying Spikes from descendant lineages were clearly more infectious than their immediate predecessors, as illustrated by comparing BA.2.75 with BA.2.75.2, XBB with XBB.1.5, XBB.1.16 with XBB.1.16.1, EG.1 with EG.5, EG.5 with EG.5.1, and BA.2.86 with JN.1; the latter evolved from the heavily mutated BA.2.86 through the acquisition of only a single additional S-domain mutation. BA.2.87.1 maintains infectivity similar to the suboptimal strain BA.2.86, but approximately two times lower than the successful JN.1 circulating at the time of BA.2.87.1 emergence. These results show that epistatic effects do not fully compensate for the impact of the numerous mutations in BA.2.87.1 Spike. Nevertheless, they also highlight that antigenically distinct viruses, which evolved to escape immunity at the expense of optimal fitness, as previously observed with XBB or BA.2.86, can rapidly acquire compensatory mutations restoring small infectivity defects.

**Fig. 1.**
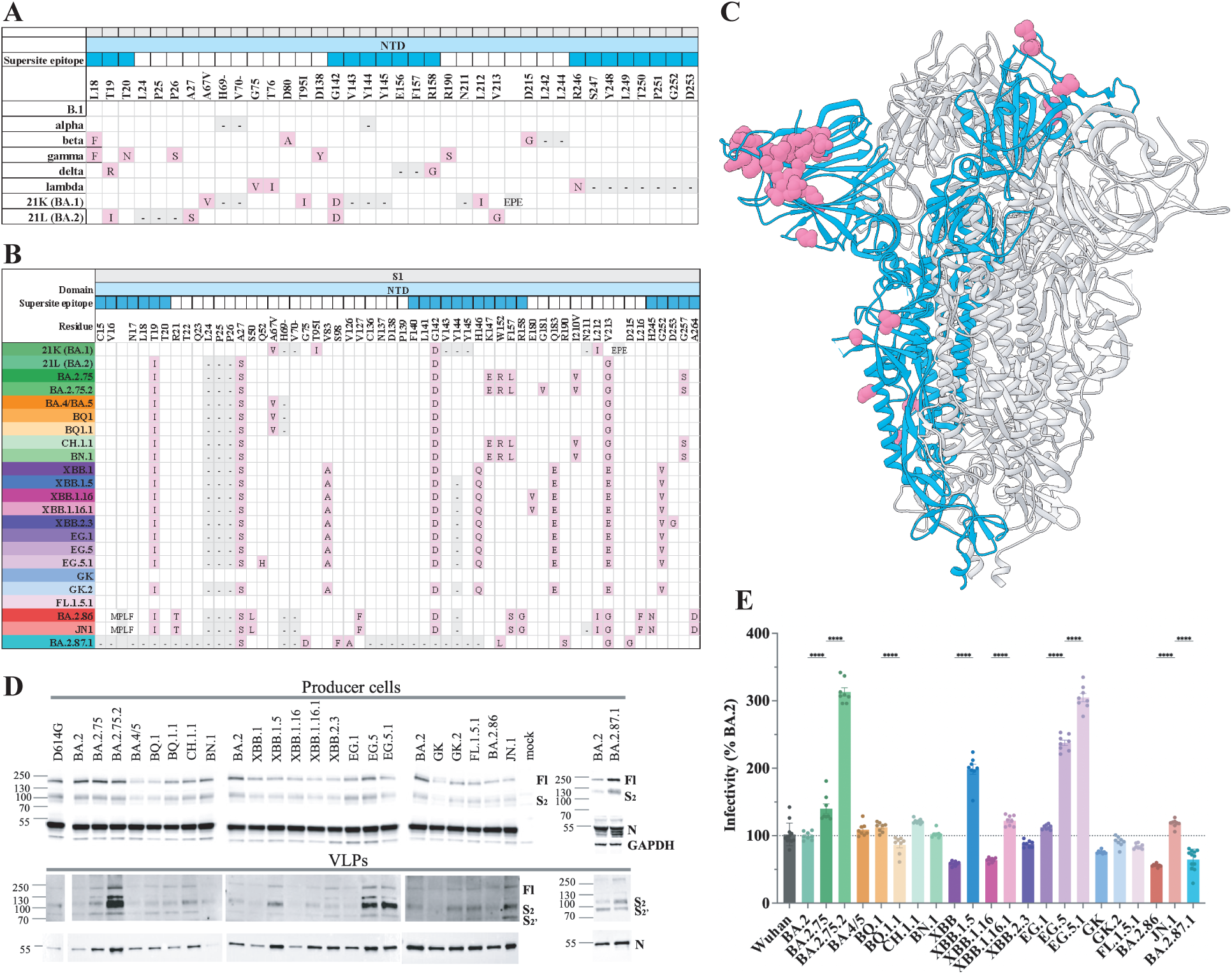
Mutations in the NTD of Spike variants and infectivity of SC2-VLPs. (**A**) The nature and position of the mutations indicated in the NTD are relative to the Wuhan Spike. Pink squares indicate mutations and grey squares insertions or deletions. Domains overlapping with the NTD antigenic supersite are indicated on top of the table in dark blue. (**B**) Same analysis as in (A) for the different Omicron sublineages. (**C**) Ribbon representation of the Wuhan Spike atomic model (PDB ID: 7JJI), with one protomer coloured in blue and residues uniquely mutated in BA.2.87.1 (absent from all the 22 other strains), highlighted in pink. (**D**) Top panel: SC2-VLPs stocks were produced in duplicates in HEK293T cells and lysates of the producer cells were loaded onto SDS-PAGE and analyzed by western blotting using anti-Spike (S) and anti-Nucleocapsid (N) antibodies. Equal loading was verified with anti-GAPDH antibody. Only one representative replicate is shown. Fl: full length Spike, S2: S2 domain of the cleaved Spike. Bottom panel: A genome equivalent amount of particles was purified on sucrose cushion, loaded onto SDS-PAGE and analyzed by western blotting using anti-S and anti-N antibodies. (**E**) SC2-VLPs stocks were used to transduce HEK293T ACE2^+^ TMPRSS2^+^ cells (n ≥ 6 technical replicates for each biological duplicate). Transduction efficiency was monitored by Luciferase assay. Results were normalized for genome content of each stock and BA.2 values set at 100%. Bars represent mean ± SD. For clarity, only selected statistical data are shown. \*\*\**P* < 0.001 (Ordinary one-way ANOVA test).

### Local structural rearrangements and greater loop flexibility in BA.2.87.1 Spike

Next, we applied cryo-EM to structurally characterize the BA.2.87.1 Spike (**Fig. 2 and fig. S3**). A stabilized version of the ectodomain was produced in HEK923F cells and purified as described in the Methods section. The Spike protein was loaded onto electron microscopy grids, and a dataset of 6’631 micrographs was collected (**Fig. 2A**). From these data, we resolved the overall trimer structure at a global map resolution of 3.26 Å (**Fig. 2B and fig. S3**). However, the dynamic nature of the NTD and RBD resulted in lower local resolution for these regions. Hence, we applied a strategy based on focused classification and local refinement to obtain a higher quality reconstruction of NTD and RBD (**Fig. 2C and. fig. S3**), enabling more precise characterization of structural changes induced by the mutations. The final global map resolution was 3.3 Å with RBD and NTD domains at a local resolution of ∼4 Å (**fig. S3**). The corresponding map and model information are provided in tables S1 and S2, respectively.

**FIG. 2.**
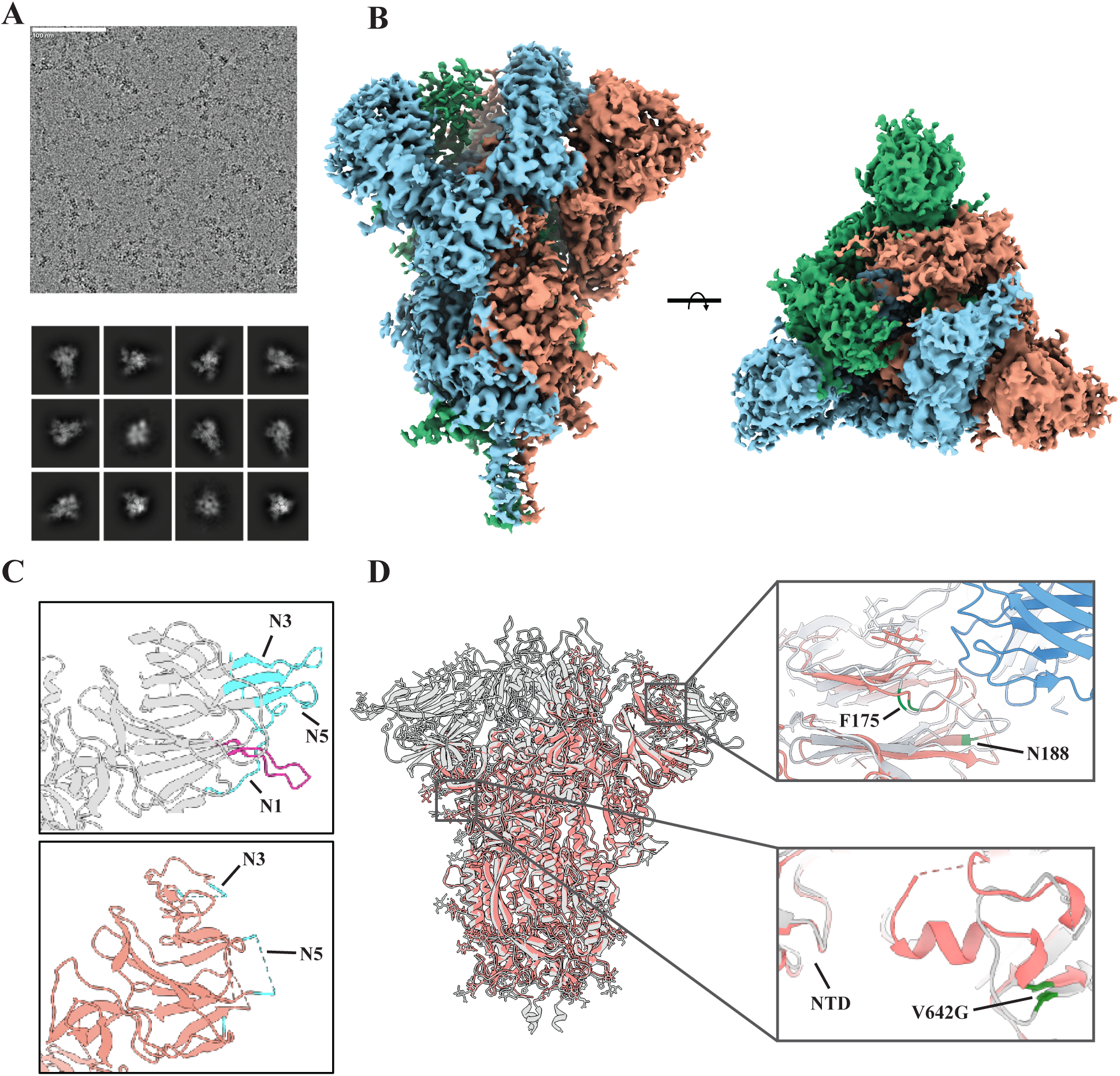
Structural analysis of BA.2.87.1 by cryo-EM. (**A**) Examples of a micrograph obtained by cryo-EM (Top) and result of the 2D classification of the particles (Bottom). (**B**) 3D map reconstruction of the trimeric Spike. Each monomer is coloured differently. Side and top views are shown. (**C**) Ribbon representations comparing the NTDs of the previously published Wuhan (top, PDB ID: 7NT9) and BA.2.87.1 Spike proteins (bottom). In blue, the 3 epitopes forming the NTD antigenic supersite. Dashed lines represent the 4 segments of flexible loops difficult to resolve confidently in BA.2.87.1 protein. In pink, the loop formed by residues A67-D80 located outside the antigenic supersite and unstructured in BA.287.1. (**D**) Ribbon representation of the overlay of the Wuhan (in grey) and the BA.2.87.1 (in orange) atomic models. The top close-up view illustrates a similar rearrangement of the NTD central loop including the F175 residue (in green), in both the BA.2.87.1 Spike and the Wuhan Spike bound by an anti-NTD antibody (in blue). Residue N188 bound by a glycan in BA.2.87.1 is highlighted in green. The bottom close-up view illustrates the conformational change imposed on the preceding loop by the V642G mutation in BA.2.87.1.

Overall, the map and model are in close agreement with the previously published structure of BA.2.87.1 Spike (*16*), and despite 65 mutations, the polypeptide backbone closely resembles the ancestral Wuhan Spike (PDB ID: 7NT9 (*15*); **Fig. 2D**). Compared to Wuhan, the NTD and RBD at the apex of the molecule exhibit greater flexibility, resulting in several poorly resolved regions within the NTD (e.g., residues L244-G261 in the N5 loop, **Fig. 2C**) and RBD (e.g., residues 410-485) that could not be modelled. Among the resolved regions, a notable change is the formation of a short helix spanning residues C617 to N641 in the S1 Domain D-loop, likely facilitated by the V642G mutation (**Fig. 2D**). Favourable packing of hydrophobic residues W633 and Y636 stabilizes this helix, which makes contacts with the NTD base. While the impact on function or stability cannot be directly inferred, this rearrangement may alter the presentation of conformational epitopes, potentially affecting antigenicity.

Large NTD deletions are a defining feature of BA.2.87.1 (**Fig. 1, B and C**). A 13-residue N-terminal deletion removes the first N1 segment of the antigenic supersite and eliminates a disulfide bond between residues C15 and C136 altogether destabilizing amino acids A67-D80 that form a loop in the Wuhan Spike. Despite 11 out of 14 identical amino-acids between the two spikes in this region, the segment is completely unstructured in BA.2.87.1 (**Fig. 2C**). A second large deletion spanning residues C136-H146 in the N3 loop partially removes the antigenic supersite and induces substantial rearrangements of the downstream 9 residues, as well as the neighbouring hydrophilic loop F175-K187, highly conserved among sarbecoviruses (*15*) (**Fig. 2D**). This loop adopts a conformation similar to that reported for biliverdin-depleted Spike bound to a monoclonal antibody (*15*). The metabolite-binding pocket collapses, with F175 occupying the volume previously filled by biliverdin in published Spike structures (e.g., PDB ID: 7NT9 (*15*)).

Within this remodeled region, the R190S substitution emerges as a key change likely contributing to pocket destabilization. In the ancestral Spike, R190 contributes to biliverdin binding through favorable cation–π interactions and salt-bridges with carboxylate moieties (**Fig. 3A**). Its replacement by a smaller, uncharged serine in BA.2.87.1 destabilizes the pocket architecture. This substitution also introduces a novel N-linked glycosylation sequon, resulting in glycan attachment at N188. Interestingly, the N188 residue is virtually invariant across the >17 million sequences deposited to GISAID (https://gisaid.org/, as of January 2026) and the entropy value below 1 for this site indicates a functional importance of this residue (**Fig. 3B**). The R190S mutation, which was previously transiently detected in the Gamma variant in November 2020, has recently raised in frequency with the emergence of newer variants such as XFG (*26*).

**FIG. 3.**
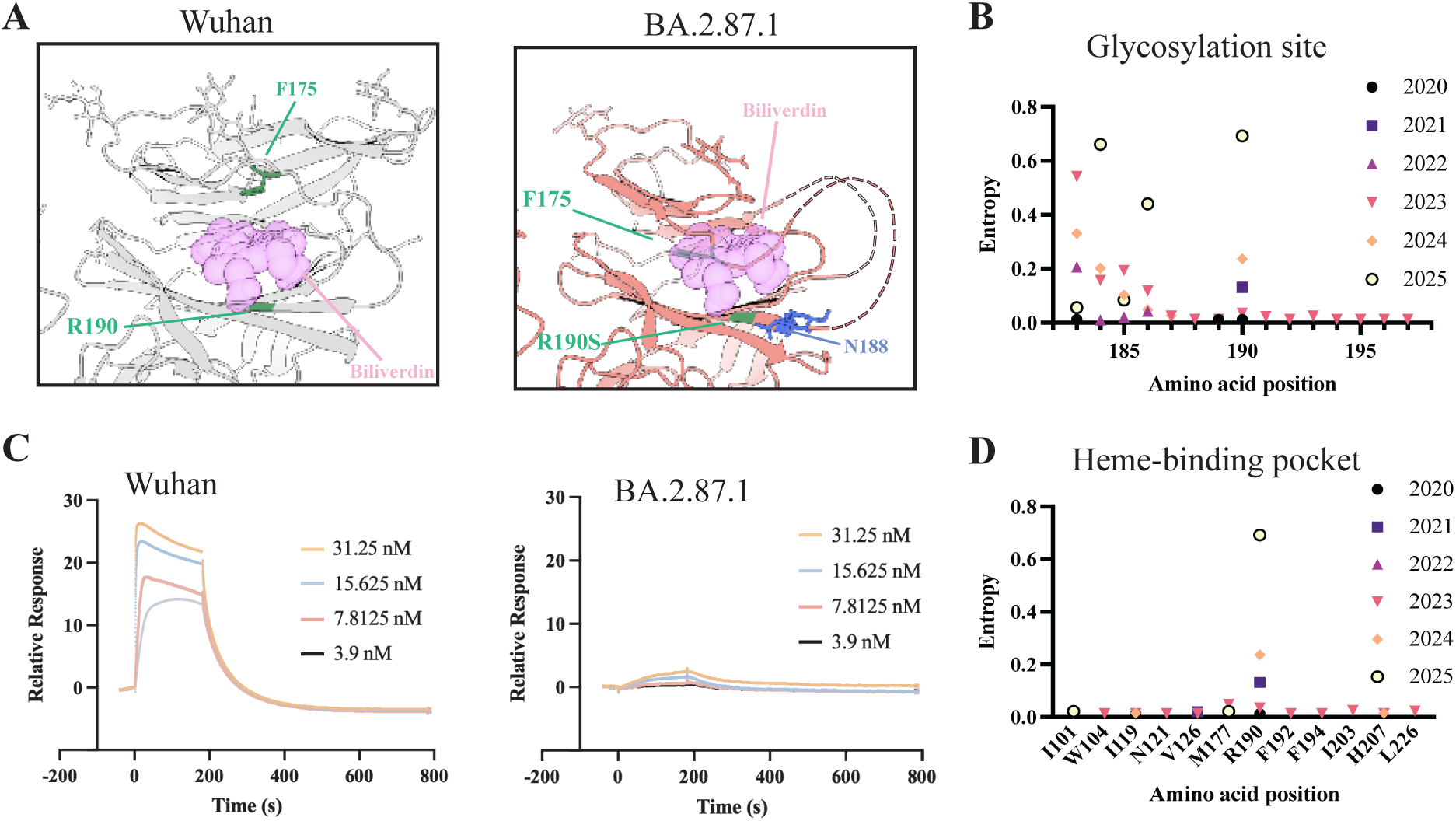
Loss of heme-binding in BA.2.87.1. (**A**) Ribbon representations comparing the heme-binding pocket of Wuhan (PDB ID: 7NT9) and BA.2.87.1. Biliverdin atoms are shown as pink spheres. F175 (green) in BA.2.87.1 occupies the space taken by the biliverdin in the Wuhan Spike. The N-linked glycan at position 188 (blue) is also partially blocking access to the space in BA.2.87.1. (**B**) Entropy values for residues surrounding the N188 glycosylation site are indicated. Values, obtained for each year since 2020 and coloured differently were extracted from Nextstrain. (**C**) SPR sensorgrams were recorded using Wuhan (Top) or BA.2.87.1 (Bottom) Spike. Purified biliverdin-depleted proteins were immobilized on a sensor chip and binding and dissociation of biliverdin was recorded at indicated concentrations. (**D**) Entropy values for residues in the heme-binding pocket are indicated. For each position, values obtained each year since 2020 were extracted from Nextstrain (https://nextstrain.org).

Altogether, analysis of the BA.2.87.1 structure and sequence indicates that extensive immune-driven evolution can reconfigure antigenic surfaces within the NTD, through coupled biochemical and conformational changes, including glycan remodeling, while preserving similar overall architecture.

### BA.2.87.1 shows very low affinity for biliverdin

Next, we focused on the absence of biliverdin-associated density in the BA.2.87.1 structure. Previous studies have shown that binding of heme metabolites such as biliverdin conditionally modulates NTD supersite conformation by altering flexible loop arrangements, thereby influencing recognition by certain classes of monoclonal antibodies (*15*). Residues found in close contacts with the heme-metabolite in the S protein structure (I101, W104, I119, N121, V126, M177, R190, F192, F194, I203, H207 and L226) are either invariant or highly conserved among sarbecoviruses, including SARS-CoV-1 (*15*). Consistently, the sequence entropy value for all contact residues is below 1 (https://gisaid.org/), confirming the functional importance of this site (**Fig. 3C**). Based on the BA.2.87.1 NTD structure, we hypothesized that mutations in the heme-metabolite lining pocket, particularly R190S, may compromise biliverdin binding. To test this, we purified the BA.2.87.1 and Wuhan Spike ectodomains using size-exclusion chromatography (SEC) under mild acidic conditions known to deplete Spike from biliverdin (*15*). Native polyacrylamide gel electrophoresis confirmed that both proteins remained trimeric after acid treatment (**fig. S4**). Biliverdin binding was then compared using surface plasmon resonance (SPR) with immobilized, biliverdin-depleted Spikes (**Fig. 3C**). The resulting dissociation constants (K_D_) for Wuhan and BA.2.87.1 were 5.8 ± 2.8 nM and 935 ± 912 nM, respectively. The large uncertainty range for BA.2.87.1 reflects a near-background binding signal, consistent with very low affinity. Given that physiological concentrations of free biliverdin and related heme metabolites are typically in the nano- to micromolar range, BA.2.87.1 Spike is therefore expected to remain largely unliganded, consistent with the structural observations (**Fig. 3A**).

The coordinated changes in loop conformations, disruption of the metabolite-binding pocket, and local glycan shielding represent major structural remodeling and are expected to collectively reshape conformational epitope presentation in this region.

### BA.2.87.1 is less susceptible to neutralization by human serum antibodies

To determine whether the structural alterations observed in BA.2.87.1 translate into changes in antigenic properties, we first examined interactions with monoclonal and polyclonal antibodies. The NTD-targeting monoclonal antibody 4A8 (*27*) was produced recombinantly, while polyclonal IgGs were purified from sera of eight convalescent donors. Enzyme-linked immunosorbent assays (ELISA) were used to assess the binding of each antibody sample to the trimeric ectodomain of BA2.87.1 and Wuhan Spikes (**Fig. 4A**). Deletions within the NTD abolished binding of 4A8, as evidenced by a near-complete loss of detectable ELISA signal (**fig. S5A**). When tested with polyclonal antibodies, BA.2.87.1 exhibited an average 2.3-fold reduction in EC_50_ values compared to Wuhan, indicating generally reduced antibody reactivity (**Fig. 4A** and **fig. S5B**).

**FIG. 4.**
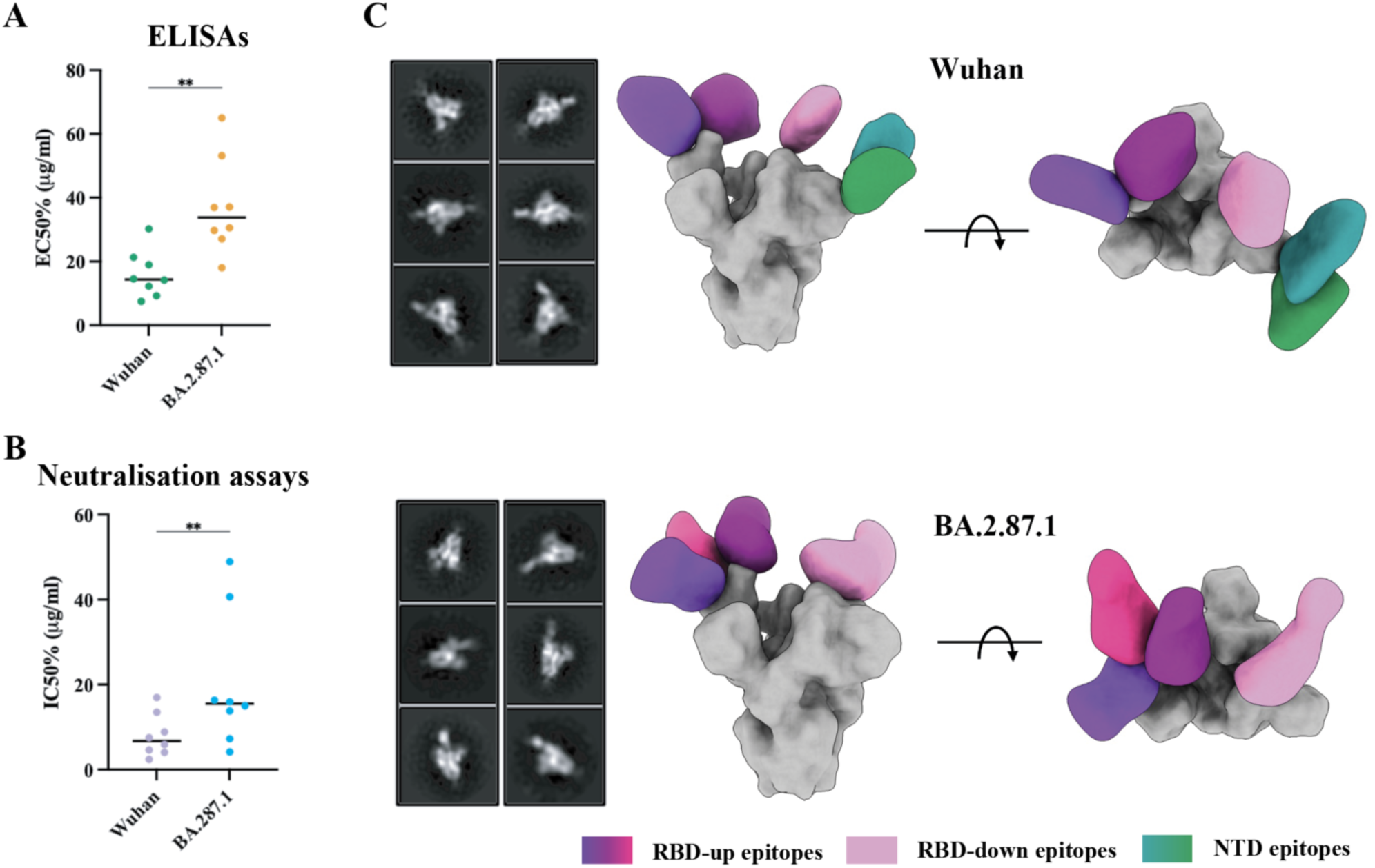
Comparison of Wuhan and BA.2.87.1 immunogenicity. (**A**) The binding of IgGs isolated from 8 different donors, to Wuhan and BA.2.87.1 Spike proteins, was quantified by ELISA. The concentration resulting in 50% of the maximal signal (EC_50_) is given for each antibody. \*\**P* < 0.01 (Wilcoxon matched-pairs test), n = 4 replicates for each antibody at each concentration point (**B**) The results of neutralization assays performed using the same set of polyclonal antibodies probing the SARS-CoV-2 VLPs pseudotyped with the Wuhan or the BA.2.87.1 Spikes. The half maximal inhibitory concentration (IC_50_) for each antibody is given. \*\**P* < 0.01 (Wilcoxon matched-pairs test), n = 4 replicates for each antibody at each concentration point. (**C**) The results of nsEMPEM analysis performed using the Wuhan (top) and BA.2.87.1 (bottom) Spikes in complex with the polyclonal Fabs from the 8 donors. In each panel, 2D class-averages are shown on the left and composite figures from the epitope mapping analysis on the right. Spike proteins are represented in grey and antibodies are coloured according to the epitope-cluster targeted.

The same panel of antibodies was then applied to evaluate the neutralization capacity against the SC2-VLPs pseudotyped with the two Spikes. Neutralization of Wuhan pseudoviruses was relatively uniform across donors with an average IC_50_ of 8 µg/ml (**Fig. 4B** and **fig. S5C**). Conversely, BA.2.87.1 pseudoviruses showed greater inter-donor variability and a higher mean IC_50_ of 20.3 µg/ml. Together, these results demonstrate reduced susceptibility of BA.2.87.1 to donor-derived antibodies, consistent with the reduced binding observed by ELISA. However, the persistence of measurable neutralization suggests that a subset of nAb epitopes remain largely intact.

### nsEMPEM confirms the loss of NTD epitope recognition by human polyclonal antibodies

Previous studies have shown that the RBD and NTD represent the principal targets of neutralizing antibodies. While mutations in the RBD are known to play a major role in immune escape, BA.2.87.1 RBD lacks some of the escape mutations recurrently found in Omicron strains like the G446S, the T478K or the F486S/V/P mutations. Overall, the BA.2.87.1 RBD shares 97% sequence similarity with the parental BA.2 RBD and 94% with the ancestral Wuhan RBD, and the observed changes in antigenic properties for BA.2.87.1 are more likely driven by the extensive remodelling of the NTD including large deletions, restructuring of central loops, loss of biliverdin binding, and gain of an N-linked glycan. To structurally characterize the primary immunogenic epitopes of BA.2.87.1 compared to the ancestral Wuhan Spike, we used the negative-stain electron-microscopy-based polyclonal epitope mapping (nsEMPEM) (**Fig. 4C and fig. S6**). This approach enables visualization of the epitope landscape targeted by polyclonal serum antibodies without requiring monoclonal antibody isolation. Fabs from the 8 human sera were pooled and cleaved into Fab and Fc fragments with papain. Digested pooled Fabs were complexed with the purified ectodomains of Wuhan and BA.2.87.1 Spikes, and immune complexes were analyzed by nsEMPEM. Using the Wuhan Spike as antigen, we resolved five structurally-distinct Fab classes bound to unique epitopes. Two classes contacted the RBD in the up-conformation (RBD-up), one bound the down-conformation (RBD-down) and two targeted the NTD. The RBD-up Fabs approached the RBD tip at two distinct angles, occupying the regions commonly associated with Class I and II nAbs (*28*) and thus likely representing neutralizing specificities. Two additional Fab classes bound the NTD at near-opposite angle of approach, both contacting regions corresponding to the antigenic supersite loops. Although the limited resolution of negative-stain maps precludes residue-level analysis, antibodies targeting this NTD supersite (e.g., 4A8, COV2-2676, COV2-2489, 4-8, 5-24) have previously been shown to mediate potent neutralization (*10*, *27*), supporting the functional relevance of these interactions.

Using the BA.2.87.1 Spike, we resolved four structurally distinct types of Fabs bound to the antigen: three engaging the RBD-up and one bound to the RBD-down conformation. Among the RBD-up Fab classes, one was uniquely observed in BA.2.87.1 complexes but not in Wuhan complexes, consistent with an antibody specificity that may have arisen in individuals previously exposed to BA.2 and related variants. Strikingly, no Fab binding to the NTD was detected, consistent with the significant structural rearrangements observed on a structural and sequence level. The absence of detectable NTD-targeting antibodies in BA.2.87.1 aligns with the reduced ELISA binding titres and provides a mechanistic explanation for the observed decrease in neutralization. Nonetheless, the remaining partial neutralization of the BA.2.87.1 suggests that protection against infection by similar variants may persist as long as the RBD recognition by pre-existing serum antibodies is sufficiently preserved.

## Discussion

In this work, we focused on BA.2.87.1 as a model to explore how extensive sequence remodeling impacts Spike structure, function, and antigenicity. Using SARS-CoV-2 virus-like particles as a surrogate for authentic virus, we found that BA.2.87.1 Spike-coated particles show partially reduced yet comparable infectivity relative to 21 major Omicron lineages and sublineages. Cryo-EM analysis revealed that the overall Spike structure is largely conserved, with four notable changes in the NTD: (1) loss of the C15-C136 disulfide bond due to C15-P26 and C136-H146 deletions, (2) increased domain flexibility, (3) incorporation of a novel glycan site at N188 and (4) remodelling of the biliverdin-binding pocket. Together with smaller NTD mutations, these changes pointed towards altered antigenicity. Using a combination of ELISA, neutralization assays, and nsEMPEM applied to human polyclonal antibody samples, we showed that neutralization with pre-existing anti-SARS-CoV-2 antibodies is decreased, with a drop of engagement of the modified NTD antigenic supersite. These findings highlight the structural flexibility and antigenic adaptability of Spike, outside the RBD, providing mechanistic insight into how Spike evolution can evade pre-existing immunity.

Our data demonstrate the remarkable structural tolerance of SARS-CoV-2 Spike to substantial deletions. While substitutions in the RBD frequently enhance ACE2 binding or promote antibody escape, deletions have recurrently appeared in the N1, N2, and N3 flexible loops of the NTD throughout the pandemic. Unlike substitutions, deletions cannot be easily reverted, producing persistent and durable changes in antigenicity. The first described NTD deletion, at positions H69-V70, arose in combination with D796H in immunocompromised patients in 2020 (*29*) and was shown to compensate for the reduced infectivity caused by this escape mutation. Similar deletions have been observed in field isolates of pangolin and bat sarbecoviruses, as well as mink-adapted (“Danish spill-back”) strains. Unlike the two other deletions in BA.2.87.1, these residues, though outside of the antigenic supersite, reside in the N2 loop. Flexible loops are generally surface-exposed and more tolerant to mutations than well-ordered and/or highly coordinated secondary-structure elements, making the NTD a privileged site for evolving antigenic novelty without disrupting receptor binding or overall Spike architecture.

Another important antigenic difference lies in the region surrounding the heme-metabolite binding pocket. Historically, this site has been conserved from SARS-CoV-1 onward (*15*). Together with the emergence of the novel N188 glycan site, these changes constitute an unprecedented antigenic diversification event that may seed a new branch of SARS-CoV-2 variants characterized by more extensive NTD remodeling. Notably, while the NTD adopts a β-sandwich fold that is broadly conserved among sarbecoviruses, the connectivity of β-strands and their relative spatial arrangement differ substantially in Spikes from other coronaviruses, including HKU1, OC43, and NL63. This structural diversity indicates that far more extensive NTD evolution is structurally permissible, although the probability and evolutionary drivers of such transitions remain difficult to predict.

While the full phylogenetic history of BA.2.87.1 is unresolved, its Spike differs by 65 residues from the ancestral Wuhan strain, and 64 residues from its contemporary JN.1 variant. Based on the observed RBD reactivity, pre-existing anti-SARS-CoV-2 antibodies were likely to provide partial protection against this specific variant. However, our antigenic analyses (i.e., ELISA, nsEMPEM, and neutralization) do not resolve the reactivity of individual clones in the polyclonal antibody pool, and most antibodies in sera exhibit limited antigenic breadth. This leaves potential for escape for future similar variants through a small number of strategically positioned mutations.

Few BA.2.87.1 sequences have been detected in the world while viral infectivity, as empirically extrapolated from *in vitro* entry levels, is comparable to BA.2.86 (Fig. 1D). In BA.2.86, acquisition of a single RBD mutation (L455S) gave rise to the emergence of JN.1 variant, which temporarily became a dominant strain and is the precursor of currently circulating strains. BA.2.87.1 does not carry this mutation, nor some of the RBD mutations that have arisen through convergent evolution across different linages, such as G446S, T478K or the recurrent F486 substitutions, indicating room for further evolution of this type of variants. Hence, the extensive NTD remodeling potential of the Spike, combined with few additional compensatory or antigenicity-altering mutations in the RBD could create a variant with both enhanced replicative fitness and significantly altered antigenic properties, potentially limiting the effectiveness of pre-existing antibodies. This emphasizes the importance of monitoring variants that combine multi-domain remodeling with “subtle” RBD changes for the design of SARS-CoV-2 vaccine boosters and public health-related strategies..

## Materials and Methods

### Cell culture

The HEK293T-AT stable cell line expressing human ACE2 (A) and TMPRSS2 (T) was generated by transduction of HEK293T cells with a pRRL-ACE2-IRES-puromycin lentivector produced as described earlier (*30*), followed by puromycin selection (1 µg/ml), then transduction with a pRRL-TMPRSS2-IRES-blasticidin lentivector, followed by puromycin (1 µg/ml) and blasticidin (5 µg/ml) selection. Cells were maintained in DMEM 10% FCS containing 0.5 µg/ml puromycin and 2.5 µg/ml blasticidin.

### Spike-expressing plasmids

HexaPro plasmid was used as backbone for trimeric Spike ectodomain production (*31*). Variant Spikes were cloned as described earlier (*32*). BA.2.87.1 was inserted into the plasmid cut by XbaI and KpnI and assembled from 3 synthetic fragments (Integrated DNA Technologies) using In-Fusion® cloning (In-Fusion® Snap Assembly Master Mix, ref 638949, Takara Bio) to give the HexaPro-BA.2.87.1 plasmid. HDM-IDT-Spike-fixK plasmid (BEI catalogue number NR-52514, obtained from J.D. Bloom, Fred Hutchinson Cancer Research Center) was used as backbone to clone all the Spikes used to produce SC2-VLPs, as described earlier (*32*). BA.2.87.1 was generated by assembly of 4 PCR-amplified fragments, using HexaPro-BA.2.87.1 as template, into the HDM-Spike-BA.2 plasmid (*32*) cut with NotI, XhoI and AgeI. All the constructs were sequence verified.

### SC2-VLPs production, titration and infectivity

#### VLPs production

SARS-CoV-2 VLPs were produced as described by Syed et al. (*23*) with modifications. Briefly, for production in T75 cm2, 20 μg CoV-2-N, 10 μg CoV-2-M-IRES-E, 1.8 μg HDM-IDT Spike-fixK and 30 μg of the Luciferase-encoding Luc-PS9 plasmid were mixed and used to transfect HEK293T with Fugene® HD reagent as recommended by the manufacturer (ref E2311, Promega). Quantities were adapted according to the surface for production in 6-well plates. Media was changed 16 hours later and VLPs containing supernatants were collected 40 hours post-transfection. Supernatants were clarified by low speed centrifugation 5 min at 500 ×g, 0.45 μm filtered and single use aliquots were stored at -80°C. VLPs were titrated and normalized for the genome content by real-time quantitative reverse transcription polymerase chain reaction (RT-qPCR) as described below.

#### SC2-VLPs genome quantification

SC2-VLPs RNA was extracted from clarified supernatants in duplicates using E.Z.N.A.® viral RNA extraction kit (Omega Bio-Tek, USA) according to the manufacturer instructions with an extra DNAse I treatment step 30 min at room temperature (RT) (ref AMPD1, Sigma-Aldrich, USA). RNA was reverse transcribed with Maxima H Minus cDNA Synthesis Master Mix ( ref M1681, ThermoFisher Scientific) as recommended by the manufacturer. Each cDNA was quantified in triplicates by RT-qPCR using PowerUp SYBR green Master Mix (ref A25741, ThermoFisher Scientific) on QuantStudio Real-Time PCR Systems (ThermoFisher Scientific) with SDS 2.4 Software. Controls without reverse-transcription step were run in parallel to ensure the absence of DNA contamination in the extracted RNAs. Primers used for VLPs genome quantification are specific for the Luciferase gene: O.Luc2(f): GTGGTGTGCAGCGAGAATAG; O.Luc2(b): CTGTTCAGCAGCTCGCGCTC.

#### SC2-VLPs Western blotting

Cell extracts from producer cells were lysed with RIPA buffer (PBS + 1% NP40 + 0.5% sodium deoxycholate + 0.1% SDS). Lysates were incubated 30 min on ice, clarified by centrifugation (18000 rpm, 4°C, 30 min) and loaded on SDS-PAGE. SC2-VLPs produced as described above were normalized for genome content and purified by centrifugation 90’, 17000 rpm, 16°C, on a 20% sucrose cushion. Pelleted particles were lysed in Tris 10 mM pH8 + EDTA 1 mM pH8 + 0.1% Triton. Samples were mixed with 4x reducing Laemmli buffer (Bio-Rad, ref 1610747), denatured for 5 min at 95°C before being loaded on SDS-PAGE gels (SurePAGE Bis-Tris 4-20% Precast gel). Gels were run using MOPS buffer (GenScript, ref M00138). Gels were transferred 7 min, 15V, to PVDF membrane using a Trans-Blot Turbo transfer system (Bio-Rad Laboratories). Membranes were probed with anti-Spike (1:2000 dilution, LS-Bio, ref C19510) and anti-Nucleocapsid antibodies (1:2000 dilution, Genetex, ref GTX135357) overnight at 4°C, followed by 1 hour incubation at RT with goat anti-rabbit-HRPO antibody (1:5000 dilution, ProteinTech, ref SA00001-2). GAPDH was used as loading control and detected with anti-GAPDH-HRPO antibody (1:2000, Invitrogen, ref MA5-15738). All membranes were developed using WesternBright ECL-spray (ref K-12049-D50, Advansta, USA) and imaged using Fusion Solo S (Vilber, Germany).

#### Infectivity assay

For infectivity assays, HEK293T-AT target cells were seeded (4.10^4^ cells/well) the day before infection in 96-well plates previously coated with poly-L-lysine (0.01% w/v solution, Sigma-Aldrich USA). The day of infection medium was replaced by 50 μl of SC2-VLP-containing supernatants produced in duplicates and each tested in n=6-8 technical replicates. Luciferase assay was performed 24 hours later using Bright-Glo Luciferase Assay System (Ref E2610, Promega). After 2 min, the lysates were transferred in flat black 96-well plates (ThermoFisher scientific) and luminescence was measured with an acquisition time = 2 sec on a Infinite Tecan microplate reader. Values for each VLP were normalized to BA.2 VLP infectivity added as control in each experiment.

#### Neutralization assay

HEK293T-AT cells were seeded (4.10^4^ cells/well) the day before infection in 96-well plates previously coated with poly-L-lysine (0.01% w/v solution, Sigma-Aldrich USA). Two-fold serial dilutions of serum samples were prepared in quadruplicates in 80 µl of DMEM 2% FCS and mixed with an equal volume of VLPs for 1 h at 37°C. The medium was removed from cells and 140 µl of the mixture was applied on cells. After 36 h, the Luciferase assay was performed using the Bright-Glo Luciferase Assay System (Ref E2610, Promega). After 2 min, the lysates were transferred in flat black 96-well plates (ThermoFisher scientific) and luminescence was measured with an acquisition time = 2 sec on an Infinite Tecan microplate reader. The average luminescence value obtained for the controls without VLPs was considered as the Max inhibition, and the average luminescence value obtained for the controls non-treated with serum was considered as the Min inhibition. The % inhibition was calculated as follows = (1- ([Test – Max inhibition] / [Min inhibition – Max inhibition]) × 100). IC_50_ values were obtained using the GraphPad Prism 10 NonLinear four parameters curve fitting analysis.

### Protein production and purification

#### Protein expression

The Wuhan and BA.2.87.1 constructs described above were used to express Spikes in transiently transfected HEK293F cells (Invitrogen, ref R70007). For the protein production, the cells were maintained in Free-Style 293 expression medium (Life Technologies) and upscaled to 1L (0.8-1.2.10^6^ cells/mL). PEI-MAX (1 mg/mL) and 500 µg of plasmid were diluted in pre-warmed Opti-MEM I Reduced Serum Medium (Life Technologies), filtered (0.2 µm) and mixed together. The mix was incubated for 30 min at RT before being added to the cells. The HEK293F transfected cells were cultured for 5 days at 37°C, in an atmosphere containing 5% CO_2_ and at a rotation speed of 130 rpm.

#### Protein purification

The SARS-CoV-2 proteins were purified from cell culture supernatants by affinity chromatography using a prepacked HisTrap HP 5 mL column (Cytiva). Briefly, the supernatants were vacuum filtered (0.2 µm) and passed through the column (with a flow rate of 2 mL/min). After column washing with PBS, bound His-tagged proteins were eluted using 10 column volume (CV) of elution buffer (PBS with a gradient ranging from 25mM to 500 mM Imidazole, pH = 7.5). The proteins were concentrated using AmiconRTM Ultra Centrifugal filters (100-kDa cut off) and buffer-exchanged in TBS (150 mM NaCl, 20 mM Tris, pH = 7.5). SARS-CoV-2 purified proteins were further fractionated by size-exclusion chromatography (SEC) using the S200 16/600 Superdex column (Cytiva) to obtain pure trimers. The trimers were concentrated using an AmiconRTM Ultra Centrifugal filter (100-kDa cut off). Protein concentrations were determined by UV_280_ absorbance using theoretical extinction coefficients obtained from Expasy (ProtParam tool).

#### Native-PAGE and SDS-PAGE

Aliquots of purified trimeric proteins were mixed with 4x NativePAGE Sample buffer (Invitrogen, ref BN2003) and loaded onto a 3-12% Bis-Tris Mini Protein gel (Invitrogen, ref BN2011BX10). The gels were run for 2 hours at 150 V using Anode-Buffer (20x NativePAGE Running Buffer, Invitrogen) and Dark Blue Cathode-Buffer (20x NativePAGE cathode Running Buffer, Invitrogen). Native-PAGE gels were destained using 30% EtOH, 10% AcOH and 60% Water. Purified Spike proteins were also analyzed by reducing SDS-PAGE. Samples were mixed with 4x Laemmli buffer (Bio-Rad, ref 1610747) supplemented with 0.1 M DTT and denatured for 5 min at 95°C, before being loaded on SDS-PAGE gels (SurePAGE Bis-Tris 4-20% Precast gel). Gels were run using MOPS buffer (GenScript, ref M00138) and stained with InstaBlue Protein Stain Solution (RayBiotech, ref 331-22405-1).

### BA.2.87.1 cryo-EM

#### Cryo-EM-grids preparation

Grids were prepared using a Vitrobot Mark IV device (ThermoFisher Scientific). The following Vitrobot settings were used: temperature = 10 °C, blotting force = 10, wait time = 5 seconds, blotting time = 3 seconds. Three microliters of BA.2.87.1 (3mg/ml) were loaded onto the plasma-cleaned ultra Au Foil R 300 grids 1.2/1.3 (Quantifoil Micro Tools, ref Q350AR13A). The grids were plunged into liquid nitrogen-cooled liquid ethane and stored in liquid nitrogen until imaging.

#### Cryo-EM data collection

Cryo-EM grids were loaded into a Glacios electron microscope (ThermoFisher Scientific) operating at 200 kV. The microscope was equipped with an X-FEG and a Falcon 4 direct electron detector camera. Exposure magnification was set to 150,000 with the resulting pixel size at the specimen plane of 0.926 Å. Automated data collection was performed using EPU (ThermoFisher Scientific), and the total electron dose was setup to 50 e^-^/Å^2^. In total 6631 micrographs were collected for this entire dataset. Specific information on the imaging settings can be found in Table S1.

#### Cryo-EM data processing

Raw EER files were aligned and dose-weighted using Patch Motion Correction in cryoSPARC Live (*33*). The output micrographs were then imported into a cryoSPARC.v4 project workspace, and subjected to Patch CTF. Spike particles were picked using template picker, and three rounds of 2D classification were applied to eliminate bad particles picks and disassembled trimers. Particle stacks comprising 633’423 Spikes were transferred to Relion/4.0 (*34*) for further processing. After one round of 3D auto refinement with C3 symmetry imposition, the particles were subjected to 3D classification with local mask around the Spike NTD to separate a subset of particles with homogenous density for this region. The classification was performed without image alignment and with the regularization parameter (T) set to 16. The classification yielded 8 classes, two of which were selected for further refinement. The 3D auto refinement was performed with a soft solvent mask around the entire Spike trimer, and with restriction of angular searches to a local range of 3.7°. The C1 symmetry was imposed for this step. The resulting half-maps were used for postprocessing producing the final map used for model building. Due to the 3D classification approach, only one NTD/RBD combination was resolved well in the map, while the other two appear as scattered density due to flexibility. Alternative approaches for processing were also tested (including symmetry expansion and partial image subtraction) but without major improvement to the map. The final map was reconstructed using 216’028 particles. The data processing workflow, 2D classes, relevant maps and statistics are illustrated in Figure S3.

#### Model building

The initial model of the BA.2.87.1 was generated using SWISS-MODEL (*35*), and docked into the postprocessed map from the previous step using UCSF Chimera (*36*). The 3D atomic model was generated using the map restraints through a combination of manual model building in Coot (*37*) and automated refinement in Rosetta (*38*). As mentioned above, the reconstructed map contained only one NTD-RBD combination resolved into homogeneous map at ∼4Å resolution, so the final model was evaluated using MolProbity (*39*) and EMRinger (*40*) metrics. The model and corresponding maps and masks were deposited to the Protein Data Bank (PDB) and Electron Microscopy Data Bank (EMDB), respectively. Model and map statistics are shown in Table S2.

### ELISA

SARS-CoV-2 trimers, diluted at 3 µg/mL in 0.1 M NaHCO_3_ buffer, were coated for 2 hours at RT on Clear Flat-Bottom 96-well ELISA plates (BD Biosciences, 353279). The plates were then blocked using PBS + 5% BSA + 0.05% Tween-20 buffer overnight at 4°C with a sealing cover. Unbound trimers were removed with 3 washing steps using TBS + 0.1% Tween-20. After the washes, 30 µl of purified IgGs from patients were added in 2-fold serial dilutions and incubated for 2 hours at RT. Unbound IgGs were removed with 3 additional washing using TBS + 0.1 % Tween-20. Goat anti-human antibody (Jackson ImmunoResearch, ref 109-055-003) was added at a 1:5000 dilution in PBS + 1% BSA and incubated for 1 hour at RT. Unbound secondary antibodies were removed with 3 final washes using TBS + 0.1% Tween-20. Colorimetric detection was performed using 1-step PNPP substrate developing solution (ThermoFisher Scientific,). Absorption at 405 nm was registered using either Bio-Rad Benchmark microplate reader or an Infinite Tecan microplate reader. Negative control wells without patients IgGs and positive control wells with anti-Spike antibody (ADG-2 from Adagio(*41*)) were included in each plate.

### Biliverdin depletion and Surface Plasmon Resonnance (SPR)

#### Acidic depletion

To remove the bound biliverdin, both Wuhan and BA.2.87.1 purified trimeric proteins were treated with 0.5 M NaAc (pH = 5.2) for 30 min at RT. The treated trimers were further purified by SEC using the S200 16/600 column in 150 mM NaCl, 20 mM Tris, pH = 7.5 buffer. The depleted trimers were concentrated using AmiconRTM Ultra Centrifugal filter (100-kDa cut off) and buffer-exchanged in TBS + 1mM EDTA. Purity and integrity of the depleted proteins were verified on Native-PAGE.

#### Surface Plasmon resonance (SPR)

The binding of biliverdin to the Wuhan and BA.2.87.1 Spikes was analyzed by SPR using the purified biliverdin-depleted Spikes. All SPR analyses were performed on a Biacore 8K instrument at 25°C, using 150 mM NaCl + 50 mM HEPES (pH = 8) as the running buffer. The depleted Spike trimers were diluted in 10 mM NaAc (pH = 4.0) to be immobilized on a CM5 sensor chip (ref 29104988, Cytiva), with a contact time of 420 s and a flow rate of 10 µL/min. To collect kinetic binding data, biliverdin concentrations ranging from 0 nM to 1000 nM in 2-fold dilutions were injected at a flow-rate of 30 µL/min and a contact time of 180 s. Data were fit to a simple 1:1 interaction model using the Multi-cycle analysis option available within Biacore^TM^ Insight software (Cytiva, Biacore 8K).

### Antibodies purification and Fabs cleavage

#### Donor serum samples

Sera from the Swiss population were purchased from the Interregional Blood Transfusion Swiss Red Cross. Antibodies are representative of local immunity in September 2023, at the time of XBB* circulation. Briefly, blood from 8 donors was spun 15 min at 3000 rpm. The supernatant was collected, inactivated at 55°C for 1 hour, filtered using 0.1μm centrifugal filters and 1% Triton-X was added.

#### IgGs purification

IgGs from the 8 sera were purified on Akta Pure system (Cytiva) using a custom-made Tricorn column (Cytiva) filled with 5ml of CaptureSelect IgG-Fc resin (ThermoFisher Scientific). Following antibody immobilization on resin, the column was washed with 8 CV of phosphate-buffered saline (PBS) buffer (pH = 7.5). The antibodies were then eluted using 0.1 M glycine pH = 3, with a gradient ranging from 0% to 100%. The samples were immediately neutralized with one quarter volume of 1 M Tris-HCl pH = 8. AmiconRTM Ultra Centrifugal filters (30-kDa cut-off) were used to concentrate the purified IgGs and buffer exchange into PBS buffer.

#### IgGs digestion into Fabs

IgGs samples from all donors were pooled together and digested as described before (*42*). IgGs (0.5 mg/mL) were digested for 5 hours at 37°C using 0.02 mg/mL of Papain in 100 mM Tris, 2 mM EDTA, 10 mM L-Cysteine (pH = 7-7.5). Non-digested IgGs were separated from Fabs by SEC using the S200 16/600 column and PBS buffer. Fab/Fc-containing fractions were concentrated to 3-5 mg/mL using AmiconRTM Ultra Centrifugal filter (10-kDa cut-off) and buffer-exchanged to TBS.

### nsEMPEM samples preparation, data collection and processing

#### Samples preparation

For complexing, Fabs from the 8 pooled donors were used. 15 µg of SARS-CoV-2 trimeric protein were incubated with 3 mg of the pooled Fabs mix overnight at room temperature. The formed complexes and the unbound Fabs were separated by SEC using the Superose 6 10/300 column (Cytiva). The complexes were concentrated using AmiconRTM Ultra Centrifugal filter (100-kDa cut-off). Protein concentrations were determined using UV_280_ absorbance and theoretical extinction coefficients obtained from Expasy (ProtParam tool). The complexes were then diluted to 20 µg/mL with TBS buffer. The samples were directly deposited onto carbon-coated 300-mesh copper grids (EMS/Electron Microscopy Sciences) and stained immediately with 2% (w/v) uranyl formate for 60 seconds. Grids were imaged at 120 KeV on BioTalos with CETA camera (ThemoFisher Scientific) at 57’000 nominal magnification with the resulting pixel size at the specimen plane of 2.44. Micrographs were collected using EPU software (ThemoFisher Scientific).

#### Data processing

Preliminary processing including particle picking and 2D classification was conducted in cryoSPARC software (*43*) and the clean particle stack was transferred to Relion/4.0 (*34*). The resulting stacks had 263’079 particles for Wuhan and 340’501 particles for BA.2.87.1. Particles were then 2D-classified into 200 classes (25 iterations), and particles from 2D classes featuring antigen-Fab complexes were selected for 3D analysis. The initial 3D classification was performed into 50 classes, allowing full angular search during particle alignment. A low-pass filtered Wuhan Spike map was used as a reference for all 3D steps. Particles from similar-looking classes were then combined and reclassified to isolate different Fab-looking density. The selected 3D maps were further processed using 3D auto-refinement, and these refined maps were submitted to EMDB. The EMDB accession IDs are provided in the Data and Materials Availability section. UCSF ChimeraX 1.10.1 was used to visualize and segment the Fab density of the 3D maps.

### Statistical Analyses

*P* values were obtained with GraphPad Prism 10 for each experiment. Details on tests used are given in the appropriate figure legends. In all analyses, \*\**P* < 0.01, \*\*\**P* < 0.001.

## Acknowledgments

We thank Florence Pojer, Kelvin Lau and Yoan Duhoo from the EPFL Protein Production and Structure Core Facility for their respective advices, help with the SPR data analyses and the cryo-EM grids preparation. We thank the teams at the Interdisciplinary Centre for Electron Microscopy (CIME) and the Dubochet Center for Imaging (DCI) in Lausanne for their assistance with collection and processing of electron microscopy data. We also want to thank Vanessa Genet and Myriam Lamrayah for help with the SC2-VLPs.

## Author contributions

Conceptualization: AA, PT

Methodology: PT, EE, CR

Investigation: PT, EE, CR

Visualization: PT, EE, AA

Resources: AA, DT

Supervision: AA

Writing—original draft: PT, AA, EE

Writing—review & editing: AA, PT, EE, DT

## Data and materials availability

EM data have been deposited to PDB and EMDB servers under accession numbers: 9TMF and EMD-56058, EMD-56059, EMD-56060, EMD-56061, EMD-56062, EMD-56063, EMD-56064, EMD-5605865, EMD-56067 and EMD-56069. All data are available in the main text or the supplementary materials. All the constructs generated are available upon request to authors.

