## Supplementary material for "Antigenic landscape of a highly mutated SARS-CoV-2 Spike in ongoing viral evolution": supplem material

### FRONT MATTER

#### Title

#### Supplementary Materials

**Tables S1 and S2**

**Figures S1-S6**

**Table S1**

| | |
|---|---|
| **Microscope** | Glacios |
| **Voltage (kV)** | 200 |
| **Detector** | Falcon 4 |
| **Recording mode** | EER |
| **Magnification** | 151188 |
| **Movie micrograph pixel size** | 0.926 |
| **EER Number of fractions** | 50 |
| **Total dose ($e^{-}/Å^{2}$)** | 50 |
| **Nominal under focus range (µm)** | 0.5 – 2.0 |
| **Number of movie micrographs** | 6631 |

**Table S2**

| | |
|---|---|
| **Number of particles used for reconstruction** | 216028 |
| **Symmetry used** | C1 |
| **Global map resolution (Å)** | 3.26 |
| **EMDB ID** | EMD-56069 |
| **PDB ID** | 9TMF |
| **Residues** | 2170 |
| **Amino-acids** | 2138 |
| **Carbohydrates** | 32 |
| **RMSD Bonds (4σ)** | 0.020 |
| **RMSD Angles (4σ)** | 1.641 |
| **Ramachandran** | |
| **Outliers (%)** | 0.00 |
| **Allowed (%)** | 2.11 |
| **Favored (%)** | 97.89 |
| **Rotamer outliers (%)** | 0.00 |
| **Clash score** | 1.59 |
| **Molprobity score** | 0.93 |
| **EMRinger score** | 4.51 |

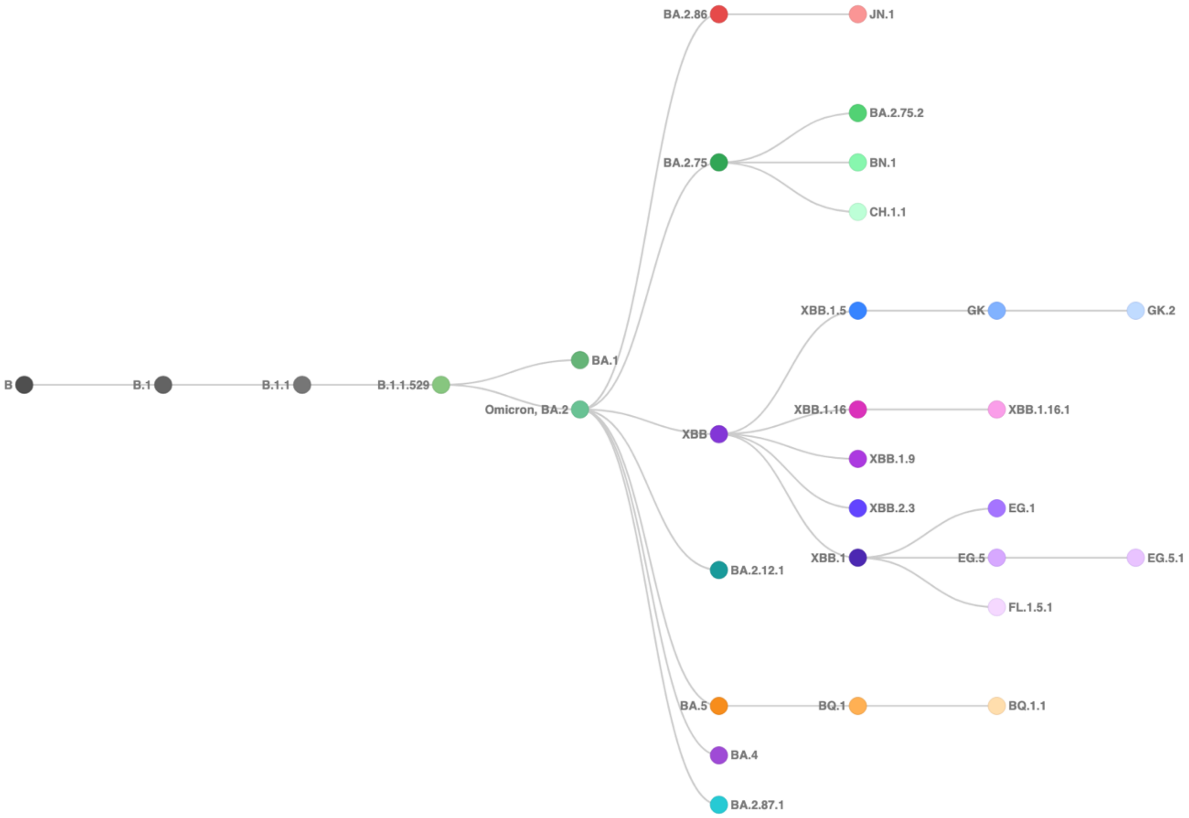


**Fig. S1. SARS-CoV-2 variants phylogenetic tree.** The SARS-CoV-2 variants phylogenetic tree is based on the full genome sequence alignment and is adapted from the NextStrain dataset (https://github.com/nextstrain/ncov-clades-schema?tab=readme-ov-file).

| | S1 | | | | | | | | | | | | | | | | | | | | | | | | | | |
|---|---|---|---|---|---|---|---|---|---|---|---|---|---|---|---|---|---|---|---|---|---|---|---|---|---|---|---|
| Domain | RBD | | | | | | | | | | | | | | | | | | | | | | | | | | |
| Supersite epitope | | | | | | | | | | | | | | | | | | | | | | | | | | | |
| Residue | I332 | G339 | R346 | K356 | L368 | S371 | S373 | S375 | T376 | R403 | D405 | R408 | K417 | N440 | K444 | V445 | G446 | N450 | L452 | L455 | F456 | N460 | S477 | T478 | N481 | V483 | E484 |
| 21K (BA.1) | | D | | | | L | P | F | | | | | N | K | | | S | | | | | | N | K | | | A |
| 21L (BA.2) | | D | | | | F | P | F | A | | N | S | N | K | | | | | | | | | N | K | | | A |
| BA.2.75 | | D | | | | F | P | F | A | | N | S | N | K | | | S | | | | | K | N | K | | | A |
| BA.2.75.2 | | D | T | | | F | P | F | A | | N | S | N | K | | | S | | | | | K | N | K | | | A |
| BA.4/BA.5 | | D | | | | F | P | F | A | | N | S | N | K | | | | | R | | | | N | K | | | A |
| BQ1 | | D | | | | F | P | F | A | | N | S | N | K | T | | | | R | | | K | N | K | | | A |
| BQ1.1 | | D | T | | | F | P | F | A | | N | S | N | K | T | | | | R | | | K | N | K | | | A |
| CH.1.1 | | D | T | | | F | P | F | A | | N | S | N | K | T | | S | | R | | | K | N | K | | | A |
| BN.1 | | D | T | T | | F | P | F | A | | N | S | N | K | | | S | | | | | K | N | K | | | A |
| XBB.1 | | H | T | | I | F | P | F | A | | N | S | N | K | | P | S | | | | | K | N | K | | | A |
| XBB.1.5 | | H | T | | I | F | P | F | A | | N | S | N | K | | P | S | | | | | K | N | K | | | A |
| XBB.1.16 | | H | T | | I | F | P | F | A | | N | S | N | K | | P | S | | | | | K | N | R | | | A |
| XBB.1.16.1 | | H | T | | I | F | P | F | A | | N | S | N | K | | P | S | | | | | K | N | R | | | A |
| XBB.2.3 | | H | T | | I | F | P | F | A | | N | S | N | K | | P | S | | | | | K | N | K | | | A |
| EG.1 | | H | T | | I | F | P | F | A | | N | S | N | K | | P | S | | | | | K | N | K | | | A |
| EG.5 | | H | T | | I | F | P | F | A | | N | S | N | K | | P | S | | | | L | K | N | K | | | A |
| EG.5.1 | | H | T | | I | F | P | F | A | | N | S | N | K | | P | S | | | | L | K | N | K | | | A |
| GK | | | | | | | | | | | | | | | | | | | | | | | | | | | |
| GK.2 | | H | T | | I | F | P | F | A | | N | S | N | K | | P | S | | | F | L | K | N | K | | | A |
| FL.1.5.1 | | | | | | | | | | | | | | | | | | | | | | | | | | | |
| BA.2.86 | V | H | | T | | F | P | F | A | K | N | S | N | K | | H | S | D | W | | | K | N | K | K | - | K |
| JN1 | V | H | | T | | F | P | F | A | K | N | S | N | K | | H | S | D | W | S | | K | N | K | K | - | K |
| BA.2.87.1 | | D | | | | F | P | F | A | | N | S | N | K | N | G | | | M | | | K | N | | K | | A |

| | S1 | | | | | | | | | | | | | | | | | | | | | S2 | | | | | | | | | | | |
|---|---|---|---|---|---|---|---|---|---|---|---|---|---|---|---|---|---|---|---|---|---|---|---|---|---|---|---|---|---|---|---|---|---|
| Domain | RBD | | | | | | | | | SD1 | | | SD2 | | | | | | | | | | | FP | | | HR1 | | | | | | HR2 |
| Supersite epitope | | | | | | | | | | | | | | | | | | | | | | | | | | | | | | | | | |
| Residue | F486 | F490 | Q493 | G496 | Q498 | N501 | Y505 | V511 | P521 | T547 | E554 | A570 | Q613 | D614 | P621 | V642 | H655 | I670 | N679 | P681 | R682 | S691 | N764 | T791 | D796 | N856 | D936 | S939 | Q954 | N969 | L981 | P1143 | D1199 |
| 21K (BA.1) | | | R | S | R | Y | H | | | K | | | | G | | | Y | | K | H | | | K | | Y | K | | | H | K | F | | |
| 21L (BA.2) | | | R | | R | Y | H | | | | | | | G | | | Y | | K | H | | | K | | Y | | | | H | K | | | |
| BA.2.75 | | | | | R | Y | H | | | | | | | G | | | Y | | K | H | | | K | | Y | | | | H | K | | | |
| BA.2.75.2 | S | | | | R | Y | H | | | | | | | G | | | Y | | K | H | | | K | | Y | | | | H | K | | | N |
| BA.4/BA.5 | V | | | | R | Y | H | | | | | | | G | | | Y | | K | H | | | K | | Y | | | | H | K | | | |
| BQ1 | V | | | | R | Y | H | | | | | | | G | | | Y | | K | H | | | K | | Y | | | | H | K | | | |
| BQ1.1 | V | | | | R | Y | H | | | | | | | G | | | Y | | K | H | | | K | | Y | | | | H | K | | | |
| CH.1.1 | S | | | | R | Y | H | | | | | | | G | | | Y | | K | H | | | K | | Y | | | | H | K | | | |
| BN.1 | | S | | | R | Y | H | | | | | | | G | | | Y | | K | H | | | K | | Y | | | | H | K | | | |
| XBB.1 | S | S | | | R | Y | H | | | | | | | G | | | Y | | K | H | | | K | | Y | | | | H | K | | | |
| XBB.1.5 | P | S | | | R | Y | H | | | | | | | G | | | Y | | K | H | | | K | | Y | | | | H | K | | | |
| XBB.1.16 | P | S | | | R | Y | H | | | | | | | G | | | Y | | K | H | | | K | | Y | | | | H | K | | | |
| XBB.1.16.1 | P | S | | | R | Y | H | | | I | | | | G | | | Y | | K | H | | | K | | Y | | | | H | K | | | |
| XBB.2.3 | P | S | | | R | Y | H | | S | | | | | G | | | Y | | K | H | | | K | | Y | | | | H | K | | | |
| EG.1 | P | S | | | R | Y | H | | | | | | H | G | | | Y | | K | H | | | K | | Y | | | | H | K | | | |
| EG.5 | P | S | | | R | Y | H | | | | | | | G | | | Y | | K | H | | | K | | Y | | | | H | K | | | |
| EG.5.1 | P | S | | | R | Y | H | | | | | | | G | | | Y | | K | H | | | K | | Y | | | | H | K | | | |
| GK | | | | | | | | | | | | | | | | | | | | | | | | | | | | | | | | | |
| GK.2 | P | S | | | R | Y | H | I | | | | | | G | | | Y | | K | H | | | K | | Y | | | | H | K | | | |
| FL.1.5.1 | | | | | | | | | | | | | | | | | | | | | | | | | | | | | | | | | |
| BA.2.86 | P | | | | R | Y | H | | | | K | V | | G | S | | Y | V | K | R | | | K | | Y | | | F | H | K | | L | |
| JN1 | P | | | | R | Y | H | | | | K | V | | G | S | | Y | V | K | R | | | K | | Y | | | F | H | K | | L | |
| BA.2.87.1 | | | | | R | Y | H | | | | | | | G | S | G | Y | | K | H | | P | K | I | H | | G | | H | K | | | |

**Fig. S2. Mutations in Omicron Spikes.** The nature and position of Spike mutations (except for the NTD shown in Fig.1B) in the indicated omicron sublineages are shown in pink and are relative to the Wuhan protein. Domains of the Spike are indicated on top of the table.

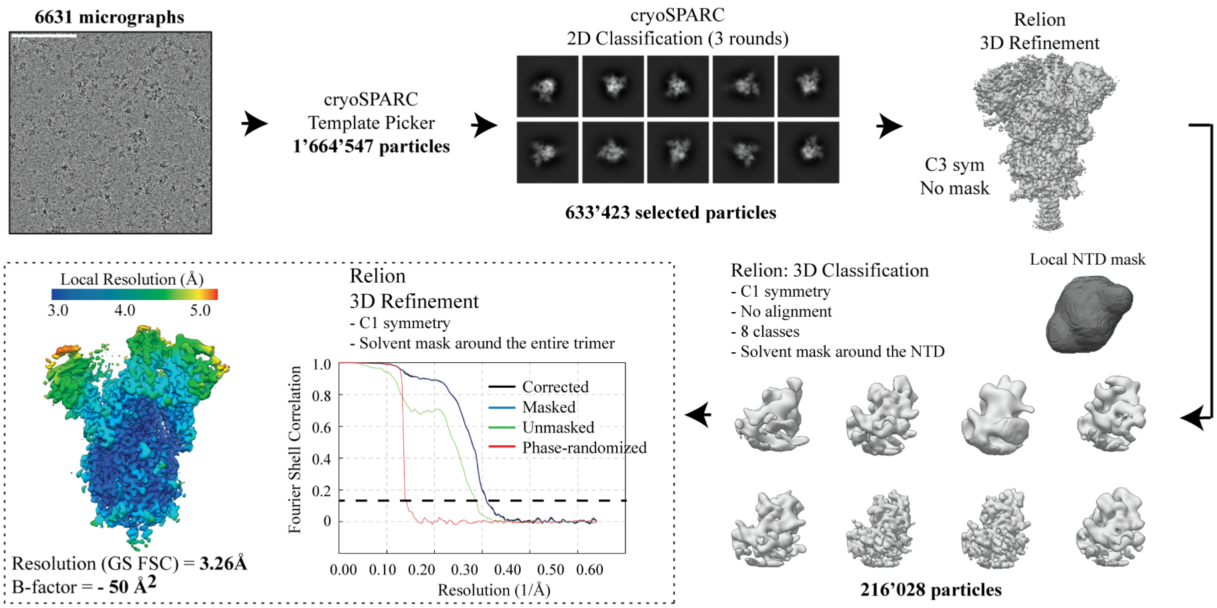


**Figure S3**. **Illustration of the data processing workflow employed to reconstruct a cryo-EM map of the BA.2.87.1 Spike**. The relevant data is shown.

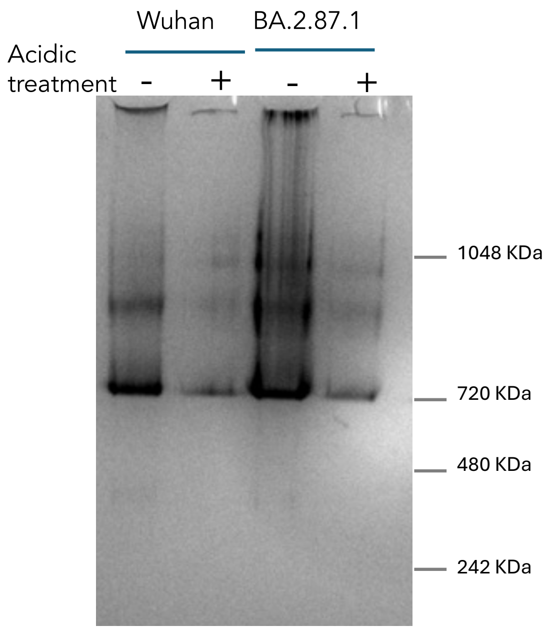


**Fig. S4. Native PAGE gels of the Spikes treated with acid.** Native gel electrophoresis of the trimeric Wuhan and BA.2.87.1 Spikes before and after treatment with low pH. The band distribution is equivalent indicating no disruption of the trimeric state.

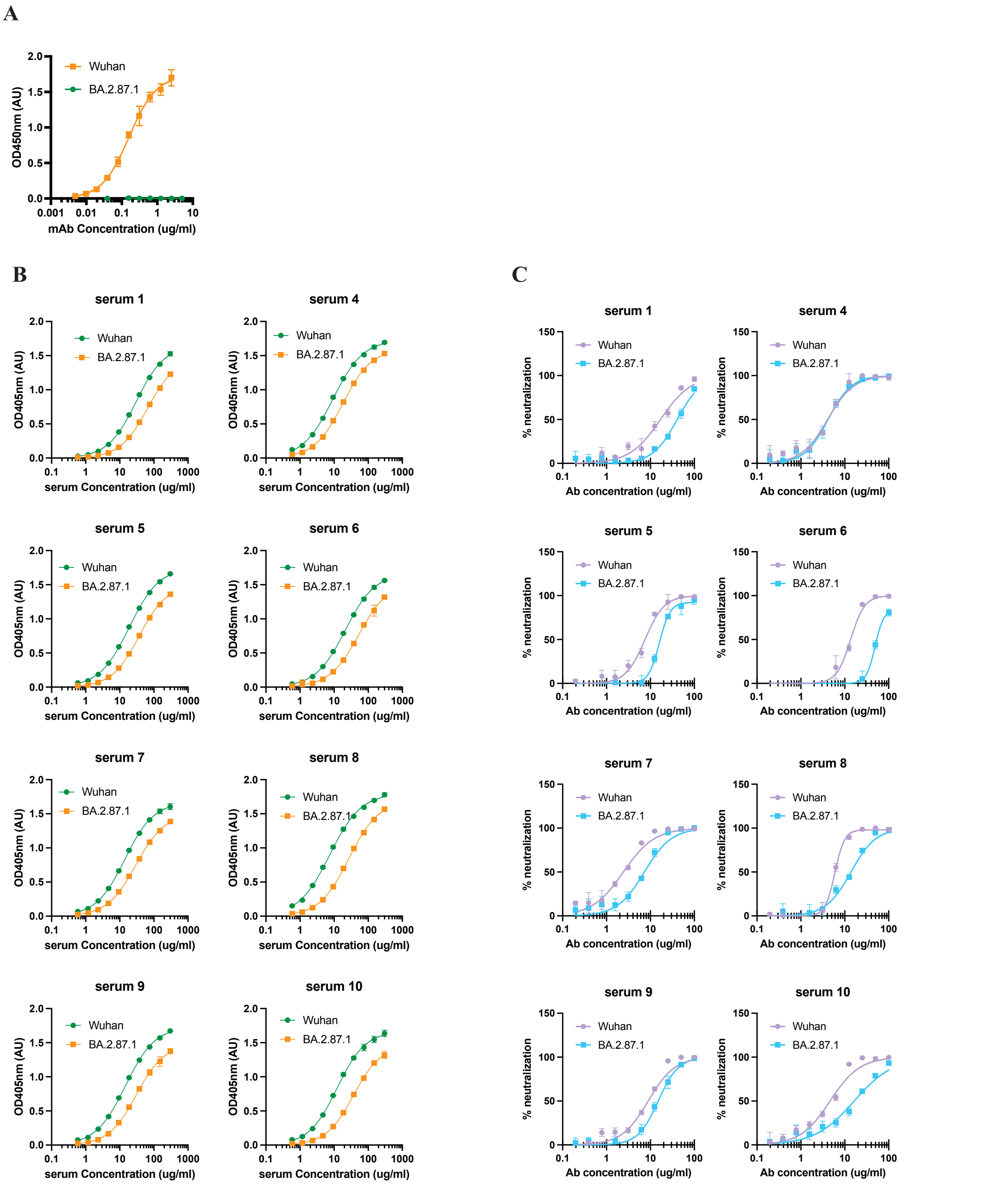


**Fig. S5. Binding and neutralization of Spikes by human antibodies.** (**A**) The BA.2.87.1 protein is not recognized by the NTD-targeting antibody as confirmed by ELISA. (**B**) The binding of IgGs isolated from 8 donors in the population was also tested by ELISA as described in Fig. 4A. (**C**) Neutralization assays were performed with the same IgGs and as described in Fig. 4B.

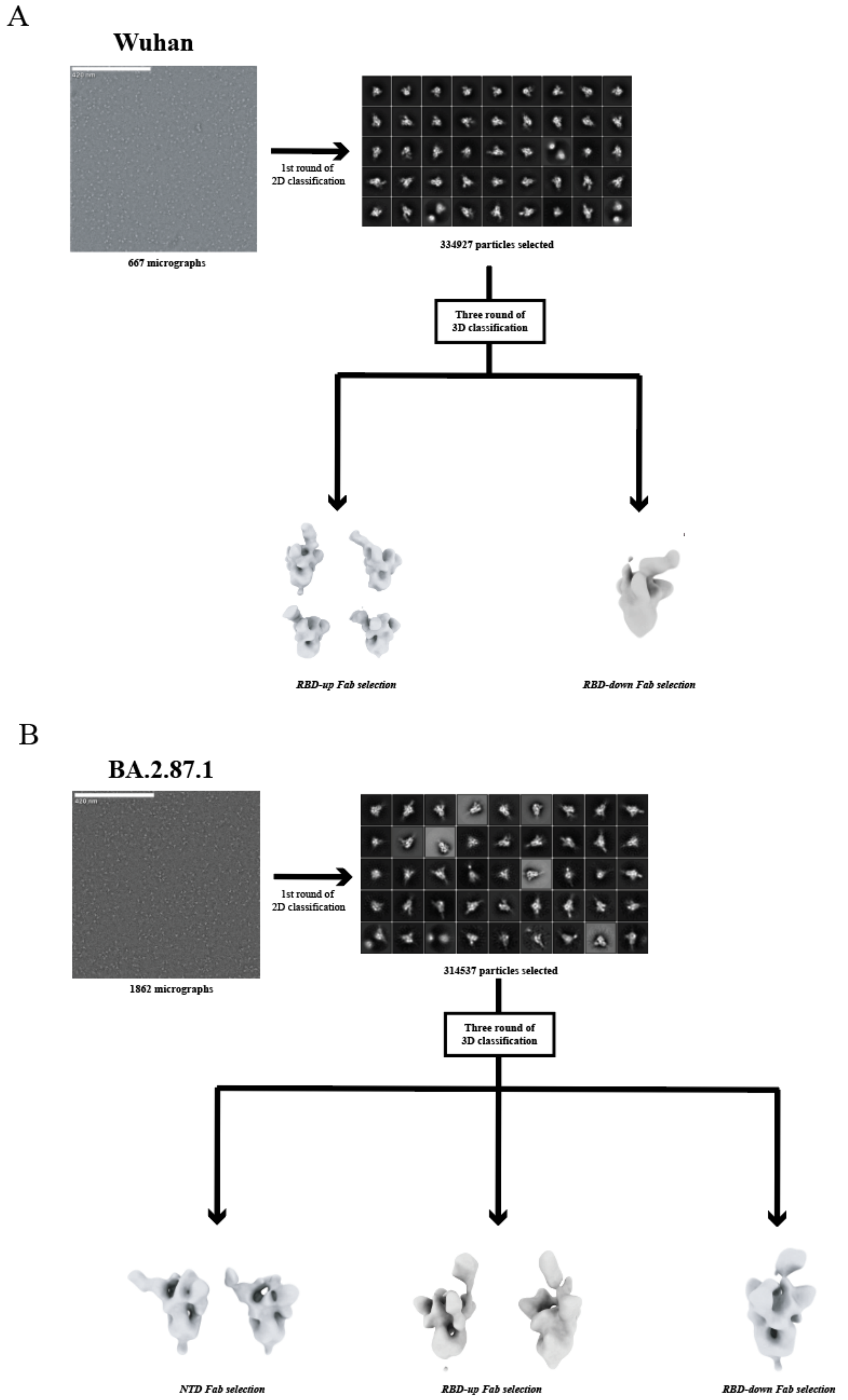


**Fig. S6. nsEMPEM data collection and processing.** The micrographs collected for the Wuhan (**A**) or the BA.2.87.1 (**B**) Spike proteins complexed with the donor Fabs were analyzed using a combination of cryoSPARC (*43*) and RELION/4.0 (*44*). A representative raw micrograph, 2D class averages and the resulting 3D classes used for Fab segmenting are shown.
